# Progranulin deficiency perturbs lipid metabolism in white matter microglia

**DOI:** 10.64898/2026.07.27.740601

**Authors:** Dahyun Yu, Ellen Armour, Sonnet Davis, Jung Suh, Matthew Simon, Jimei Tong, Gilbert Di Paolo, Leonard Petrucelli

## Abstract

Progranulin (PGRN) deficiency is a common hallmark in frontotemporal dementia (FTD) patients with granulin (*GRN*) mutations (FTD-*GRN*). Previous studies by our group and others have observed that reduced PGRN perturbs lysosomal function and microglial activation, which is believed to accelerate neurodegeneration in FTD-*GRN* patients. Lysosomal function is intrinsically linked to lipid metabolism, and evidence suggests that *GRN* deficiency can alter lipid profiles in the brain. Unfortunately, studies to date have focused on whole brain or cortical extracts, limiting our ability to assess cell type-dependent changes in lipid metabolism under disease conditions. Here, we employed lipidomic analysis specifically within the pontine microglia of *Grn* knock-out (KO) mice, a cell population that was previously linked to disease phenotypes in this model. We observed a significant reduction in the endolysosomal lipid bis(monoacylglycero)phosphate (BMP) and the lipid metabolite phosphatidylethanolamine (PE); these microglial-specific lipid alterations mirror previous whole-brain findings, suggesting that similar changes may occur across multiple cell types in the brain. We also detected a significant increase in the myelin-composing factor galactosylceramide (GalCer), which may reflect an aberrant accumulation of myelin debris within microglia that arises due to defective lysosomal clearance. Notably, lipid perturbations were exacerbated with age within *Grn* KO microglia, suggesting that changes in lipid metabolism are both age- and genotype-dependent in this model. Together, our results support our hypothesis that PGRN acts as a master regulator of critical microglial processes – including lysosomal function, lipid metabolism, and the regulation of myelination – in an age-dependent manner.

## Introduction

Frontotemporal dementia (FTD) is an umbrella term encompassing multiple neurodegenerative dementias hallmarked by atrophy of the frontal or temporal lobes, cognitive deficits, changes in behavior, and loss of executive function (1). FTD is divided into three main subgroups based on clinical presentation: behavioral variant FTD (bvFTD), non-fluent primary progressive aphasia (nfvPPA), and semantic variant primary progressive aphasia (svPPA) (2). While nfPPA and svPPA are often sporadic disorders, bvFTD has a strong genetic component: about half of bvFTD cases can be traced to a genetic cause, one of the most common being mutations in the gene granulin (*GRN*).

*GRN* encodes progranulin (PGRN), a growth factor involved in cell proliferation, inflammation, and wound healing. The protein is processed in the lysosome by cathepsin L (3), forming small proteins termed granulins, and plays a critical role in regulating lysosomal function (4-6). Disease-associated *GRN* mutations introduce a premature termination codon into its mRNA transcripts, facilitating their degradation by nonsense mediated decay (7) and suppressing PGRN expression. Accordingly, individuals with homozygous *GRN* mutations develop the lysosomal storage disorder neuronal ceroid lipofuscinosis (NCL), characterized by enlarged lysosomes and an accumulation of lipofuscin that leads to widespread progressive neuronal dysfunction (9). *GRN* haploinsufficiency, however, is strongly linked to FTD (termed FTD-*GRN*) (8), where downstream defects in lysosomal function are linked to microglial activation and cortical neurodegeneration (4-6). Approximately half of all individuals with FTD develop cytoplasmic inclusions of TAR DNA binding protein 43 (TDP-43) (10) – and this proteinopathy is strikingly more prevalent in individuals with *GRN* haploinsufficiency (11). White matter atrophy is also a common hallmark of FTD-*GRN*, although the underlying mechanisms causing this phenomenon are still poorly understood (12). White matter is enriched in oligodendrocytes (13), which are essential for the production and maintenance of myelin (14), strongly implying a link between white matter defects and synaptic dysfunction under disease conditions. Importantly, many *GRN*-associated disease hallmarks – including impaired synaptic plasticity (15), white matter atrophy (6), microglial activation (16), neuronal loss (5), increased autophagy (17), lysosomal dysfunction (4-6), and TDP-43 proteinopathy (6) – are recapitulated in homozygous *Grn* knockout (KO) mouse models, making these animals an excellent model to study the molecular mechanisms underlying FTD-*GRN*.

Lysosomal function is essential for proper lipid homeostasis, as these organelles play a central role in lipid metabolism and recycling. Lysosomal storage disorders like NCL are often marked by acute lipid accumulation within defective lysosomes, while analogous changes in lysosomal function could lead to altered brain lipidomic profiles downstream of *GRN* haploinsufficiency in FTD-*GRN* patients. In fact, previous studies in *Grn* KO mice have revealed that a full or partial loss of PGRN expression can trigger changes in the expression of endolysosomal lipids like bis(monoacylglycero)phosphate (BMP) and glucosylsphingosine (GlcSph) (18), as well as critical neuronal phosphatidylserines and energy storage lipids like triacylglycerides, diacylglycerides, and cholesterol esters (19). However, these studies were limited to lipid profiling of *Grn* KO mouse whole brain extracts or embryonic fibroblasts, hindering efforts to understand regional and cell type-dependent changes in lipid composition and processing. Previous work from our lab revealed that white matter atrophy and myelination defects in *Grn* KO mice may be due to the aberrant accumulation of myelin debris within microglial lysosomes, a phenomenon particularly evident in the pons (6). Here, we performed lipidomic analysis exclusively in pontine microglia to evaluate whether changes in lipid metabolism within this cellular population could contribute to disease phenotypes in *Grn* KO mice. Indeed, we observed unique pontine microglial lipid profiles in 12- and 21-month-old *Grn* KO mice compared to age-matched wild-type (WT) controls. *Grn* KO profiles were marked by a significant reduction in endolysosomal BMPs and cell membrane phosphatidylethanolamines (PEs) alongside a significant increase in myelin-building galactosylceramides (GalCers). Pathway analyses of differentially expressed lipids revealed an enrichment in factors associated with cellular membrane structure and function, the endoplasmic reticulum (ER), mitochondria, and the endolysosomal system in *Grn* KO pontine microglia. Together, these findings mirror previously reported changes in whole brain lipidomic profiles in *Grn* KO mice, while further uncovering specific changes in lipid regulation that occur downstream of PGRN loss in white matter microglia. Our work supports a model in which defects in lysosomal function within pontine glial cells impair myelin processing in *GRN* deficiency disorders, contributing to white matter pathology and accelerating disease.

## Materials and methods

### Mouse husbandry

All procedures involving rodents were performed in accordance with the National Institutes of Health Guide for Care and Use of Experimental Animals and approved by the Mayo Clinic Institutional Animal Care and Use Committee (IACUC; Protocol number A34315-15-R20). Mice were maintained on a 12-hour light/dark cycle in standard housing at Mayo Clinic Jacksonville with access to standard mouse chow and water *ad libitum*. All mice were of the same genetic background (C57BL/6J; The Jackson Lab #000664; RRID: IMSR_JAX:000664) and underwent the same procedures in the same order. The health and welfare of all animals were monitored daily by on-site staff. Whenever possible, individual litters were randomly divided among experimental groups to prevent any litter-specific confounding effects. Roughly equal numbers of males and females were included in each experimental cohort, and no differences between sexes were observed except for expected differences in total body weight. Mice were aged to specific time points as indicated in the text, methods, and figure legends and euthanized according to methods approved by the American Veterinary Medical Association. No animals were excluded from the study.

### Microglia isolation

Microglia were isolated from *Grn* KO and WT mice by fluorescence-activated cell sorting (FACS) using modified versions of manufacturer-established protocols. Briefly, to prepare a single cell suspension, mice were anesthetized with sodium ketamine and xylazine (90–120 mg/kg) and transcardially perfused with PBS for 5 minutes at 2.25 mL/min until all blood was flushed from the system. The pons was regionally dissected from each animal and pooled samples containing between 5 and 9 pontine tissue specimens from mice with the same sex and genotype were processed into a single cell suspension using an adult brain dissociation kit (Miltenyi Biotec 130-107-677) according to the manufacturer’s protocol. Samples were processed with myelin removal beads II (Miltenyi Biotec 130-096-731) following the debris removal step to clean up excess myelin debris, again according to the manufacturer’s protocol. Collected unlabeled cells were subsequently pelleted and resuspended in 500 µl FACS buffer (PBS/1% BSA + 1 mM EDTA) to undergo cell counting. A subset of cells was saved for individual staining controls, and both control and experimental samples were stained on ice for flow cytometric analysis with SYTOX green (1 drop per 1×106 cells) to exclude dead cells, CD11b-BV421 (BD Biosciences 562605, 1:100), and CD45-APC (BD Biosciences 559864, 1:100). Following staining, cells were pelleted and resuspended in 400 µL (experimental samples) or 200 µL (controls) FACS buffer.

Cells were strained through a 100 mm filter before sorting CD11b+/CD45Intermediate microglia on a FACS Aria III (BD Biosciences) with a 100 mm nozzle. Sorted cells were collected in a PBS/0.5% BSA solution, pelleted, and flash-frozen for future liquid chromatography-mass spectrometry (LCMS) and lipidomic analyses.

### Liquid chromatography-mass spectrometry (LCMS)

FACS-sorted microglia were extracted with LC-MS-grade 400 µL methanol containing internal standards, adjusted to 800 µL with LC-MS-grade water, and mixed with 800 µL MTBE. Samples were vortexed and centrifuged at 14,000 × g for 10 min at 4°C to induce phase separation. The organic phase was transferred to a glass vial, and a 50 µL aliquot was reserved for GlcCer, GalCer, GlcSph, and GalSph analysis (see separate method section). The remaining organic phase was dried under a stream of N2 and reconstituted in LC-MS-grade methanol for nonpolar lipid analysis.

### Lipidomic analysis

Lipid analyses were performed by liquid chromatography (UHPLC Nexera X2, UHPLC ExionLC) coupled to electrospray mass spectrometry (QTRAP 6500+). For each analysis, 5 μL of sample was injected on a BEH C18 1.7 μm, 2.1 ×1 00 mm column (Waters) using a flow rate of 0.25 mL/min at 55°C. For positive ionization mode, mobile phase A consisted of 60:40 acetonitrile/water (v/v) with 10 mM ammonium formate + 0.1% formic acid; mobile phase B consisted of 90:10 isopropyl alcohol/acetonitrile (v/v) with 10 mM ammonium formate + 0.1% formic acid. For negative ionization mode, mobile phase A consisted of 60:40 acetonitrile/water (v/v) with 10 mM ammonium acetate + 0.1% acetic acid; mobile phase B consisted of 90:10 isopropyl alcohol/acetonitrile (v/v) with 10 mM ammonium acetate + 0.1% acetic acid. The gradient was programmed as follows: 0.0-8.0 min from 45% B to 99% B, 8.0-9.0 min at 99% B, 9.0-9.1 min to 45% B, and 9.1-10.0 min at 45% B. Electrospray ionization was performed in positive or negative ion mode. For the QTRAP 6500+, we applied the following settings: curtain gas at 30 psi (negative mode) and curtain gas at 40 psi (positive mode); collision gas was set at medium; ion spray voltage at 5500 V (positive mode) or -4500 V (negative mode); temperature at 250°C (positive mode) or 600°C (negative mode); ion source Gas 1 at 55 psi; ion source Gas 2 at 60 psi; entrance potential at 10 V (positive mode) or -10 V (negative mode); and collision cell exit potential at 12.5 V (positive mode) or -15.0 V (negative mode). Data acquisition was performed using Analyst 1.6.3 (Sciex) in multiple reaction monitoring mode (MRM). Area ratios of endogenous metabolites and surrogate internal standards were quantified using MultiQuant 3.02 (Sciex).

### Analysis of bis(monoacylglycero)phosphate (BMP)

BMP analyses were performed by liquid chromatography (UHPLC Nexera X2, UHPLC ExionLC) coupled to electrospray mass spectrometry (QTRAP 6500+). For each analysis, 5 μL of sample was injected on a BEH C18 1.7 μm, 2.1 ×1 00 mm column (Waters) using a flow rate of 0.35 mL/min at 62°C. Mobile phase A consisted of 60:40 acetonitrile/water (v/v) with 10 mM ammonium acetate; mobile phase B consisted of 90:10 isopropyl alcohol/acetonitrile (v/v) with 10 mM ammonium acetate. The gradient was programmed as follows: 0.01–3.0 min from 50% B to 90% B, 3.0–3.01 min to 50% B, and 3.01–3.50 min at 50% B. Electrospray ionization was performed in negative ion mode applying the following settings: curtain gas at 30 psi; collision gas at medium; ion spray voltage at 4500 V; temperature at 600°C; ion source Gas 1 at 50 psi; ion source Gas 2 at 60 psi. LCMS data acquisition was performed using Analyst 1.6.3 (Sciex) in MRM. BMP species were quantified using a surrogate internal standard of BMP(14:0/14:0). Quantification was performed using MultiQuant 3.02 (Sciex).

### Absolute quantification of BMP(22:6/22:6) and BMP(18:1/18:1)

To quantify absolute levels of BMP(22:6/22:6) and BMP(18:1/18:1) in biological matrices (i.e., tissue lysate), aliquots of calibration standards (STDs), quality control samples (QCs), and study samples were transferred to a clean 96-well plate and extracted with methanol containing BMP(14:0/14:0) as an internal standard. After mixing and centrifugation, the supernatant was further diluted as needed and injected into LCMS for analysis. LCMS analyses were performed on an ExionLC AD ultra-high-performance liquid chromatography (UHPLC) system coupled with a Sciex API 6500+ Triple Quad mass spectrometer (AB Sciex, Redwood City, CA). HPLC was performed using an Acquity BEH C8 column (1.7 mm, 5032.1 mm; Waters Co., Milford, MA), and the column was kept at 55°C during the run. For the LC separation of BMP species of interest from the matrix interferences, two mobile phases were used: 0.01% ammonium hydroxide, 1 mM ammonium acetate in water (Mobile Phase A) and 0.01% ammonium hydroxide in acetonitrile (Mobile Phase B). The combined flow rate was kept at 0.4 mL/min. The reference standards and internal standard were purchased from Avanti Polar Lipids (Alabaster, Al). To prepare a calibration curve and assess QCs, standard stock solutions were spiked in surrogate matrices to avoid the MS signal contributions from endogenous BMP species in authentic biological matrices. During each run, a matrix QC prepared from pooled authentic biological matrices was also used to monitor the assay performance.

### Analysis of GlcCer, GalCer, GlcSph, and GalSph

Glucosylceramide, galactosylceramide and glucosylsphingosine analyses were performed by liquid chromatography (UHPLC Nexera X2, UHPLC ExionLC) coupled to electrospray mass spectrometry (TQ 6495C). For each analysis, 8 μL of sample was injected on a HALO HILIC 2.0 μm, 3.0 × 150 mm column (Advanced Materials Technology, PN 91813-701) using a flow rate of 0.48mL/min at 45 °C. Mobile phase A consisted of 92.5/5/2.5 ACN/IPA/H2O with 5 mM ammonium formate and 0.5% formic acid. Mobile phase B consisted of 92.5/5/2.5 H2O/IPA/ACN with 5 mM ammonium formate and 0.5% formic acid. The gradient was programmed as follows: 0.0–2 min at 100% B, 2.1 min at 95% B, 4.5 min at 85% B, hold to 6.0 min at 85% B, drop to 0% B at 6.1 min and hold to 7.0 min, ramp back to 100% at 7.1 min and hold to 8.5 min. Electrospray ionization was performed in positive mode. Agilent TQ 6495C was operated with the following settings: gas temp at 180°C; gas flow 17 L/min; nebulizer 35 psi; sheath gas temp 350°C; sheath gas flow 10 L/min; capillary 3500 V; nozzle voltage 500 V. Data acquisition was performed using Analyst 1.6 (Sciex) in MRM. Lipids were quantified using a mixture of stable isotopically labeled internal standards. Quantification was performed using MultiQuant 3.02 (Sciex).

### Statistical analysis

All analytes were run in technical replicates of 2–3. Studies with multiple assays were designed to be powered for the most variable analyte based on previous data. No animals were excluded from the study. Statistical comparisons for sorted cell lipidomic heatmaps were performed using MetaboAnalyst 4.0 (https://www.metaboanalyst.ca/). All heatmaps were generated in R using the Complex Heatmap package. Whenever possible, investigators were blinded to the sex and genotype of experimental samples.

## Results

### The lipidome of white matter microglia is profoundly altered in PGRN-deficient mice

To better understand the implications of white matter atrophy in *GRN*-mediated disease progression, we conducted lipidomic analyses in white matter microglia from homozygous *Grn* KO mice. Specifically, we extracted pontine microglia from 12- and 21-month-old *Grn* KO mice, as we previously determined that these timepoints recapitulate early and late-stage *Grn* deficiency disease features (6), and compared the resultant lipid profiles to those of age-matched controls. We observed significant changes in the levels of several lipid classes including PEs, BMPs, and GalCers (Fig 1A). PEs are key components of cell membranes that play significant roles in regulating lipid metabolism, dysregulation of which is implicated in several diseases ranging from neurodegeneration to liver disease (20). We observed a significant reduction in PEs in *Grn* KO mice compared to WT mice (Fig 1B), suggesting disrupted lipid processing and metabolism that could contribute to alterations in energy metabolism, cell signaling, and other cellular processes. BMPs have significant roles in modulating lysosomal function and are widely implicated in lysosomal storage disorders (21). We observed a significant reduction in BMPs in *Grn* KO mice compared to WT mice (Fig 1C), supporting our hypothesis that PRGN loss triggers lysosomal dysfunction within pontine microglia. Finally, GalCers are key building blocks of neuronal myelin produced by oligodendrocytes (22). We observed a significant increase in GalCers within the pontine microglia of *Grn* KO mice compared to WT mice (Fig 1D), which may reflect the aberrant accumulation of myelin debris within microglia and imply altered myelin regulation in *Grn* KO animals. Taken together, these results reveal that the lipid profile of microglia from *Grn* KO mice differs significantly from that of control mice, and the changes observed are indicative of perturbations in lysosomal function, lipid processing, and myelination. Further, similar lipidomic changes were observed at both the 12- and 21-month timepoints (Fig 1A), suggesting defects in lipid modulation occur early under *Grn* deficiency conditions and are maintained or exacerbated with age.

**Fig 1.**
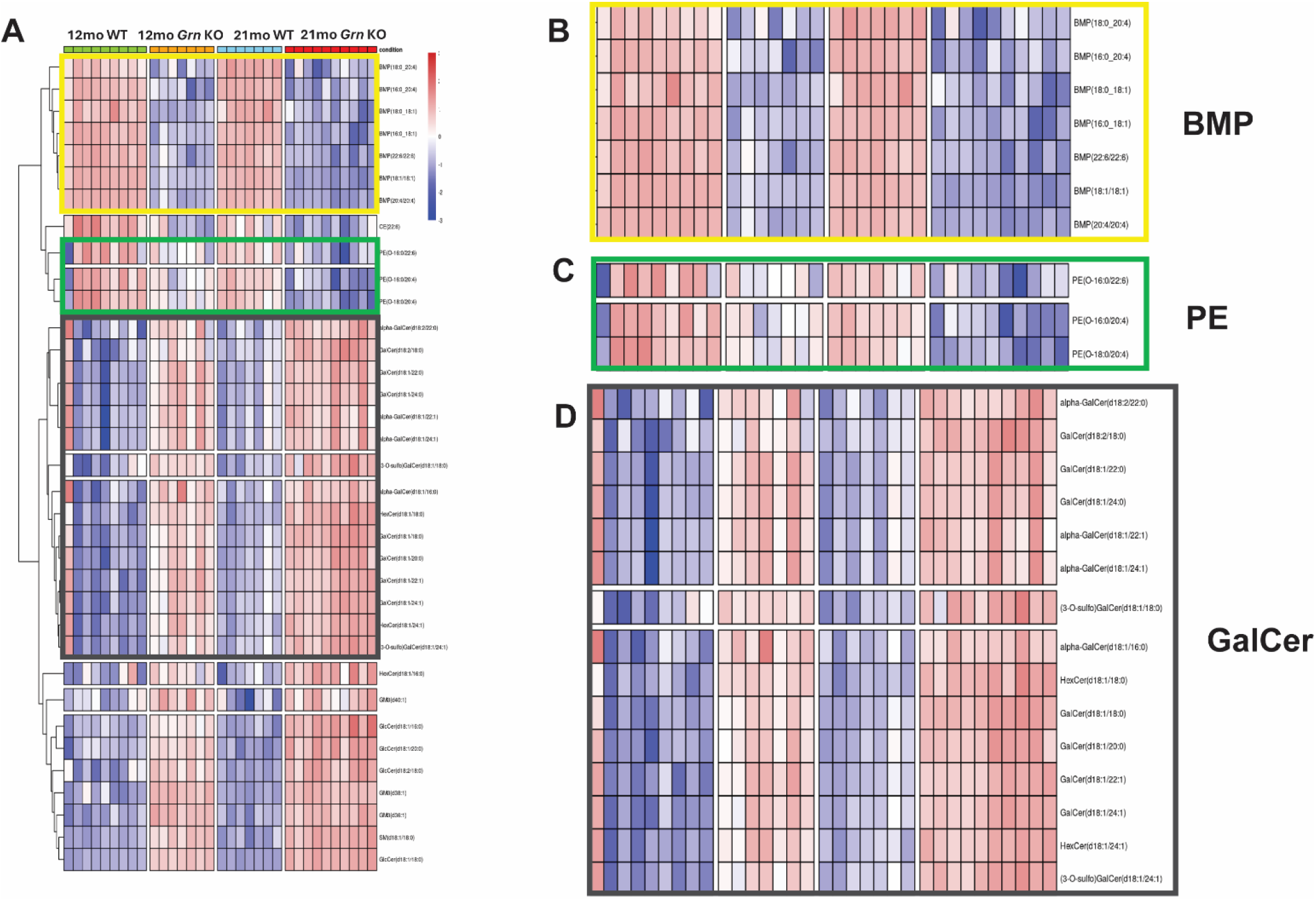
The lipidome of white matter microglia is profoundly altered in PGRN-deficient mice. (**A**) Heatmap of differentially regulated lipid species from 12- and 21-month-old *Grn* KO pontine microglia compared to age-matched WT controls. Each lane represents a sample pooled from 5–6 mouse pontine tissues. *n* = 18–25 mice/sex, *n* = 36–49 mice/experimental group, and *N* = 7–9 total pooled samples. BMP: bis(monoacylglycero)phosphate, CE: cholesterol ester, PE: phosphatidylethanolamine, GalCer: galactosylceramide, (3-O-sulfo)GalCer: sulfatide, HexCer: hexosylceramide, GlcCer: glucosylceramide, GM3: monosialodihexosylganglioside, SM: sphingomyelin. (**B**) Enlargement of BMP lipid panel shows significant reduction in BMP levels in *Grn* KO mice. (**C**) Enlargement of PE lipid panel shows significant reduction in PE levels in *Grn* KO mice. (**D**) Enlargement of GalCer lipid panel shows significant increase in GalCer levels in *Grn* KO mice.

### Endolysosomal and myelin-enriched lipids are dysregulated in PGRN-deficient pontine microglia, and defects are exacerbated with age

Given that the lipidome of *Grn* KO mice differs significantly from that of WT mice (Fig 1), we next generated volcano plots to separate and visualize individual differentially expressed lipid groups. These plots emphasized a significant reduction in BMPs in both 12- and 21-month-old *Grn* KO mice compared to WT controls (Fig 2A, B). This reduction was more evident at the 12-month timepoint, suggesting that changes in BMP levels – and by extension, lysosomal disruption – occur early in *Grn*-associated disease progression. We again observed a significant upregulation in GalCers (Fig 2A, B), which may be evidence of myelin dysregulation in *Grn* KO mice. This upregulation was apparent at both timepoints, which could suggest that myelin debris accumulation in pontine microglia is a persistent defect or that GalCer level changes stabilize with age (Fig 2B). Overall, the 21-month-old mice exhibited more elevated levels of differentially expressed lipids in comparison to the 12-month-old mice. Indeed, age-genotype analyses confirmed that changes in the lipidome occur in an age-dependent manner in the *Grn* KO microglia (Fig 2C).

**Fig 2.**
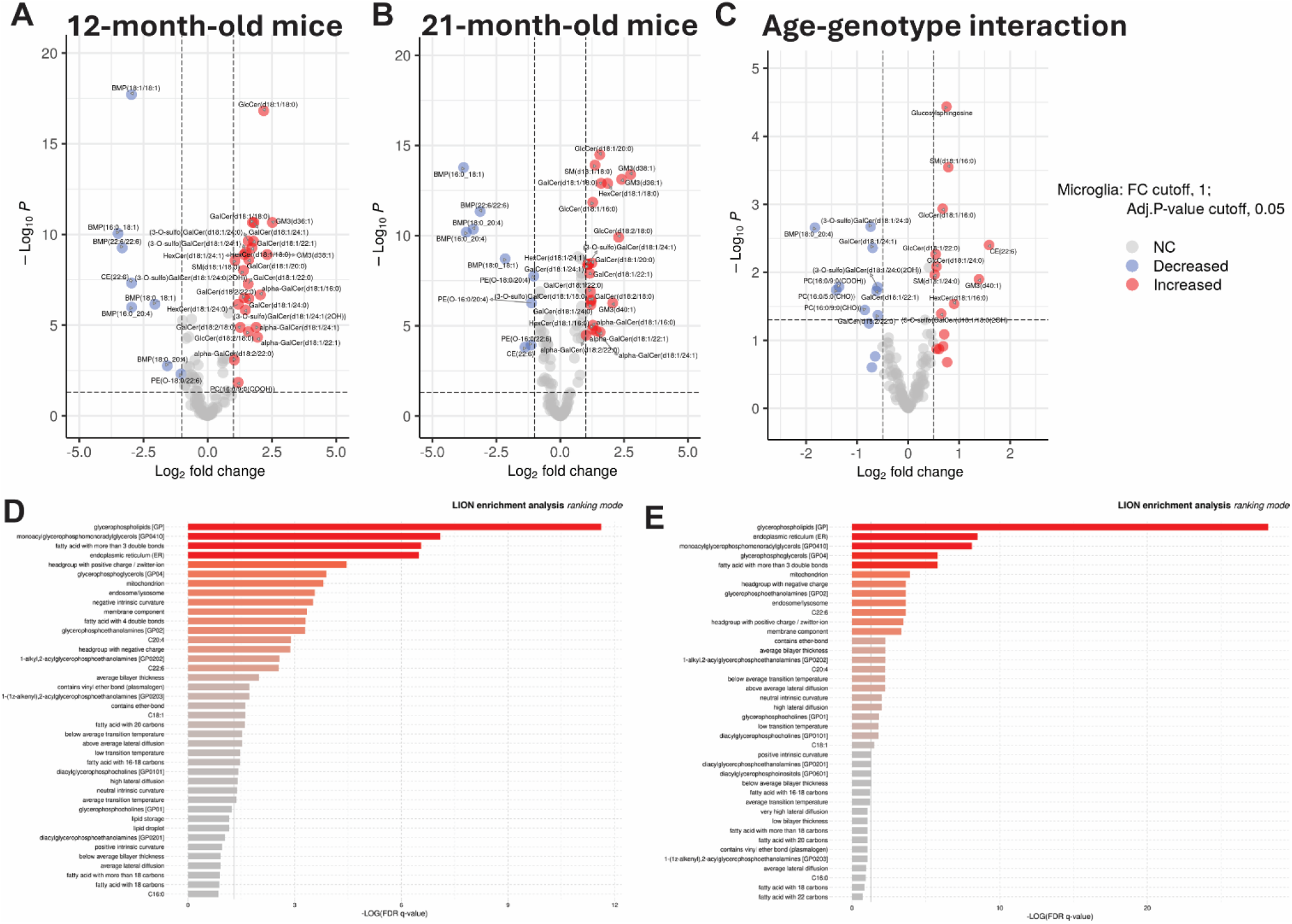
Endolysosomal and myelin-enriched lipids are dysregulated in PGRN-deficient pontine microglia, and defects are exacerbated with age. (**A–B**) Volcano plots of differentially regulated lipids in 12- (**A**) and 21-month-old (**B**) *Grn* KO pontine microglia compared to their WT counterparts. PC: phosphatidylcholine. (**C**) Volcano plot showing age-associated lipidomic changes in pontine microglia from *Grn* KO mice compared to WT controls. FC cutoff = 1, adj. *p*-value cutoff = 0.05. (**D–E**) LION enrichment analyses show differentially expressed lipids at the 12-(**D**) and 21-month (**E**) timepoints.

We next applied Lipid Ontology (LION) enrichment analyses to gain insight into the functional impact of lipid changes in *Grn* KO microglia. This process identified specific lipid groups modulated in *Grn* KO mice at each timepoint, with both age groups showing changes in glycerophospholipids and lipids associated with the ER, mitochondria, and lysosomal function (Fig 2D, E). These results imply that PGRN is essential for critical processes like protein processing and cell membrane structure and function. Notably, lipidome changes observed in the *Grn* KO pontine microglia mirror aspects of lipid profiles observed in the whole brain (18), suggesting that PGRN loss may induce similar lipid modulations across multiple cellular populations in the brain.

## Discussion

Microglia play a major role in maintaining brain function, composing approximately 5–15% of all cells in the central nervous system (CNS) (23). Their diversity contributes to this role, with microglia participating in a wide range of cellular processes such as synaptic maintenance and phagocytosis.

Microglia also mediate the inflammatory response via their activation. Therefore, it is unsurprising that microglia are implicated in many neurodegenerative diseases and metabolic disorders (24). Microglia activation is more prevalent in white matter brain regions, where it is also enriched with age (25), and microglial dysfunction impairs several neuronal processes that can ultimately lead to white matter atrophy (26). We previously observed increased microgliosis and microglial lysosomal dysfunction in *Grn* KO mice, particularly in the white matter regions of the pons (6), further supporting a role for microglia in modulating the function and morphology of the aging brain. We built upon these observations in the present study, uncovering microglial-specific lipidomic alterations in the *Grn* KO pons.

Previous studies have performed lipidomic analyses in *Grn* KO mice at varying timepoints (2–13 months of age), but these studies examined the lipid profile of the whole mouse brain (18, 19), limiting our understanding of region- and cell type-specific changes in lipid metabolism and processing. Our work helps bridge this important knowledge gap by examining lipidomic changes specifically in the pontine microglia – a cell population previously linked to disease-associated phenotypes in *Grn* KO mice (6). Moreover, we extended our analyses to 21-month-old animals, providing the first insight into lipid changes in geriatric *Grn* KO mice that are nearing disease endpoints. Strikingly, both our study and others found similar lipid modulations upon *Grn* deficiency, namely a decrease in BMPs and a concomitant increase in GlcSphs (18, 19). Alongside a reduction in BMPs, we also observed a significant decrease in PEs; downregulation of these two critical lipid groups suggests that *Grn* deficient mice are exhibiting significant lysosomal dysfunction in addition to lipid dysregulation, consistent with several previous findings indicating that loss of PGRN impairs lysosomal activity (2, 5, 6, 18, 19).

Interestingly, we also observed a significant increase in GalCers, which are one of two major classes of glycosphingolipids and key components of myelin. These findings may appear at odds with other studies, as, although there is some variation in the results, proteomic analyses have generally reported a significant reduction in the white matter-enriched myelin basic protein (MBP) in *Grn* KO mouse brain (5), and one group reported a reduction in myelin lipids in FTD-*GRN* patients (2). These results were obtained from full brain or cortical extracts, however, and we believe this net decrease in myelin components is therefore the result of white matter atrophy. The increase in myelin lipids we observed in pontine microglia is instead likely reflective of the abnormal accumulation of myelin debris within these cells arising from inefficient debris clearance within lysosomes under *Grn* KO conditions. Indeed, our results align with our previous study demonstrating lysosomal dysfunction in *Grn* KO microglia, as we reported an accumulation of myelin debris within the pons of both *Grn* deficient mice and FTD-*GRN* patients (6). Notably, another study also reported an accumulation of myelin-enriched lipids, such as sulfatide, GlcCer, and GalCer, in microglia isolated from *Grn* KO brain (18). Previous results in *Grn* KO mice indicate a significant reduction in oligodendrocyte proteins (5), as well, suggesting that these animals may exhibit a loss of oligodendrocytes with white matter atrophy. In fact, evidence suggests *Grn* KO mice fail to undergo remyelination following cuprizone-mediated demyelination (16), suggesting widespread changes in oligodendrocyte function. Together, our findings, in parallel with previous studies, support a role for PGRN in modulating lysosomal activity, lipid metabolism, and oligodendrocyte-mediated myelination. Moreover, our work suggests that lipid changes occur in an age-dependent manner in the *Grn* deficient brain, as alterations become more pronounced in animals of an advanced age.

There is still much to be learned about how *Grn* deficiency results in gliosis and neurodegeneration, but analysis of lipidomic changes in the CNS provides key insight into potential pathomechanisms. PGRN is a lysosomal protein, and the connection between *Grn* deficiency and lysosomal dysregulation is well-documented. Accordingly, a reduction in BMPs has emerged as a characteristic phenotype associated with PGRN loss in both our study and others, and this defect may underlie changes in lysosomal activity and debris clearance in affected cells (18). Mounting evidence also suggests a prominent association between lysosomal dysfunction and lipid metabolism, which has been recently linked to the pathophysiological mechanisms of neurodegeneration (32). Altered brain lipidomic profiles in FTD-*GRN* mouse models and patient samples confirm that PGRN loss (and associated changes in lysosomal activity) triggers profound changes in lipid expression that could contribute to disease outcomes.

Furthermore, while there are many signaling molecules and processes involved in myelin formation, maintenance, and remyelination, many of these factors are modulated by lipid metabolism (29) and could consequently be altered under PGRN deficient conditions. Indeed, our work links lysosomal dysfunction in *Grn* KO microglia with the aberrant clearance of myelin debris within white matter regions (6). Given the strong connection between disease phenotypes and PGRN loss, recent studies have focused on restoring PGRN expression as a therapeutic avenue for FTD-*GRN*. Results are encouraging: restoring PGRN levels corrected the lipid dysregulation observed in *Grn* KO mice (30, 31), and overexpressing even just individual granulins was sufficient to revert the lipid profile (30). Our results further highlight pontine microglia as a prospective therapeutic target. Reestablishing normal lipid profiles in this vulnerable cell population could restore lysosomal activity and the downstream clearance of myelin debris, which could then help alleviate white matter atrophy, improving oligodendrocyte-mediated myelination and neuronal function.

## Data Availability Statement

All data described in this study are included in the manuscript or figures. Materials, reagents, and resources generated as part of this study are available from the corresponding authors upon reasonable request and will be shared with qualified researchers according to guidelines set forth by the institutions involved.

